# Neighboring codon adjacent nucleotides have a conserved influence on mRNA decay

**DOI:** 10.1101/2025.11.17.688958

**Authors:** Reed S. Sorenson, Leslie E. Sieburth

## Abstract

**Background:** Degeneracy in the genetic code has been shown to play a major role in yeast mRNA stability; differences in optimality (charged tRNA supply vs demand) among synonymous codons were revealed as a determinant of mRNA decay rates. However, whether and how much this mechanism determines mRNA decay rates in plants is unsettled. Furthermore, whether factors other than charged tRNA abundance influence codon optimality remains unexplored.

**Results:** Here, biased codon usage in Arabidopsis correlated with mRNA decay rates. A codon-optimality model of mRNA decay rate based on codon frequencies was tested using synonymously recoded genes. mRNA decay rates of these alleles in transgenic plants was consistent with the model predictions. Arabidopsis appears to use the same mechanism for sensing low optimality codons as yeast and humans because the N-terminal sensor domain of NOT3 is conserved. However, decay rates were also affected by codon context. Neighboring codons’ adjacent nucleotides consistently shifted codon correlations with decay rate in Arabidopsis, which was also observed in other published datasets revealing an influence of codon sequence on translation that is independent of charged tRNA concentrations.

**Conclusions:** Codons frequencies explained 21% of decay rate variance, suggesting codon optimality-mediated decay is one of multiple mechanisms that determine decay rates in plants. Our study also found codon context as an additional factor that affects mRNA stability and establishes a paradigm of selection among synonymous codons decoded through wobble base pairing.

## Background

mRNA decay reduces mRNA abundance and thereby limits protein production. To maintain a steady RNA abundance, mRNA decay is balanced by similar rate of RNA production. In plants, this balancing act occurs over ∼2.5 orders of magnitude of RNA flux; most transcriptome-wide studies of mRNA *t*_1/2_s report ranges from a few minutes to >24 h [1–3]. While much is understood about how rates of mRNA production are specified, the cellular mechanisms by which stereotyped mRNA decay rates are determined is much less understood – particularly in plants.

mRNA species often decay by multiple pathways, so multiple mechanisms contribute to specifying a decay rate [4,5]. These mechanisms often depend on mRNA sequences or sequence features. These features can enhance or inhibit recruitment of enzymes of mRNA decay initiation (decapping, deadenylation, or endonucleolytic cleavage) which expose the RNA to degradation by processive exoribonucleases [6]. Sequence feature recognition can occur by direct interaction with RNA binding proteins (e.g., DST, AGOs, ECTs [7–13]) or by interaction with the translation machinery (e.g., premature termination codon, [14]). Recognition during translation can lead to co-translational RNA degradation, and this mode of degradation can be prevalent; for example, the Arabidopsis cytoplasmic 5’-to-3’ exoribonuclease XRN4 degrades 68% of its substrates co-translationally [15].

In this work, we focus on understanding the extent to which coding sequences influence mRNA stabilities in Arabidopsis, in a mechanism of decay initiation known as codon optimality-mediated decay (COMD, [16]). Codon optimality was defined by Ikemura [17]; codons for which there are many charged tRNAs could be decoded quickly and were designated optimal, while low tRNA availability causing slow decoding were designated non-optimal. Codon optimality is one of the reasons codons encoding the same amino acid (i.e., synonymous codons) have non-random usage in each genome analyzed [18], also known as codon usage bias (CUB). Codon optimality has been associated with tailoring translation speed for optimal folding of nascent polypeptides and protein output [19–21], slowing early translation elongation in coding sequences [22,23] and determining RNA decay rates [24]. In yeast, COMD is mediated by Not5 which senses slow ribosome transit and recruits the CCR4 deadenylation complex to the mRNA, and can account for up to 55% of decay rate variance [24,25] suggesting that this mechanism is widely used to initiate mRNA decay. In addition to yeast, mRNAs’ codon frequencies are also predictive of decay rate in zebrafish [26,27], Drosophila [28], humans [29–31], mouse [32], and mosquito [33]. Codon usage has been used to predict mRNA abundance over the last three decades [34–36] and, in plants, has long been recognized as an important feature for stabilizing foreign transgene mRNAs [37,38], but whether codon usage influences mRNA decay rates in Arabidopsis is unknown.

Here, we present evidence for both COMD and adjacent nucleotides of neighboring codons as important determinants of mRNA decay rates in Arabidopsis. We found that large variance in CUB between transcripts correlated with mRNA decay rates. Select codon frequencies accounted for 21% of mRNA decay rate variance within the whole transcriptome. Using modeled codon-decay associations, we recoded open reading frames (ORFs) which resulted in predictable shifts in mRNA stability, mRNA abundance, and mature protein abundance in transgenic plants. The codon optimality sensor domains of yeast Not5 and human CNOT3 were highly conserved in *Arabidopsis thaliana* (At)NOT3 suggesting a conserved mechanism of decay recruitment to slowed ribosomes. mRNAs for which codon frequencies were more accurate predictors of decay rate had characteristic features, such as longer 3’UTRs, a tendency for higher dependence on decapping-mediated degradation, and membership in particular functional categories. Unexpectedly, we found that codon correlations with RNA decay rate were not reading-frame dependent; out-of-frame triplet nucleotides also strongly correlated with decay rates. These correlations were due to a novel feature associated with mRNA decay rates - the influence of neighboring codons’ adjacent nucleotides. In other words, the identity of the pair of nucleotides composed of the wobble position nucleotide of a codon and the first nucleotide in the next codon had consistent associations with mRNA decay rates. Thus, codon context can significantly influence codon association with mRNA stability. This phenomenon was also observed in published data from other eukaryotes suggesting it reflects a conserved influence. The specific influence of four adjacent nucleotide pairs has broad implications for codon selection.

## Results

### Arabidopsis mRNAs have large variance in codon usage that correlates with decay rate

Codon optimality mediated decay is detectable by patterns of codon usage across a spectrum of RNA decay rates; in diverse species, optimal codons are more prevalent in mRNAs with low decay rates and non-optimal codons are used more in mRNAs with high decay rates [24–33]. To assess whether COMD functions in Arabidopsis, we first assessed the relationship between mRNAs’ codon usage and RNA decay rates by calculating codon usage bias (CUB, defined as codon count relative to the total number of synonymous codons) for representative isoforms of seedling-expressed genes (16,740 nuclear-encoded mRNAs, Sorenson et al., 2018a). Median codon usage for these mRNAs showed bias for most amino acids. For example, of the four Pro codons, median CUB of CCC and CCG was 9.5% and 15%, and that of CCA and CCU was 33% and 39%, respectively (Figure 1A). The exception was Lys codons AAA and AAG, which each showed median usage of 50% (Supplemental Figure 1A). CUB medians differed between synonymous codons, and codon usage in individual genes varied widely. For example, although the median CCU Pro usage was 39%, the interquartile range was from 30% to 48% indicating that half of mRNAs had more extreme usage (Figure 1A). This variation might exist because of distinct codon influences on mRNAs [16].

**Figure 1.**
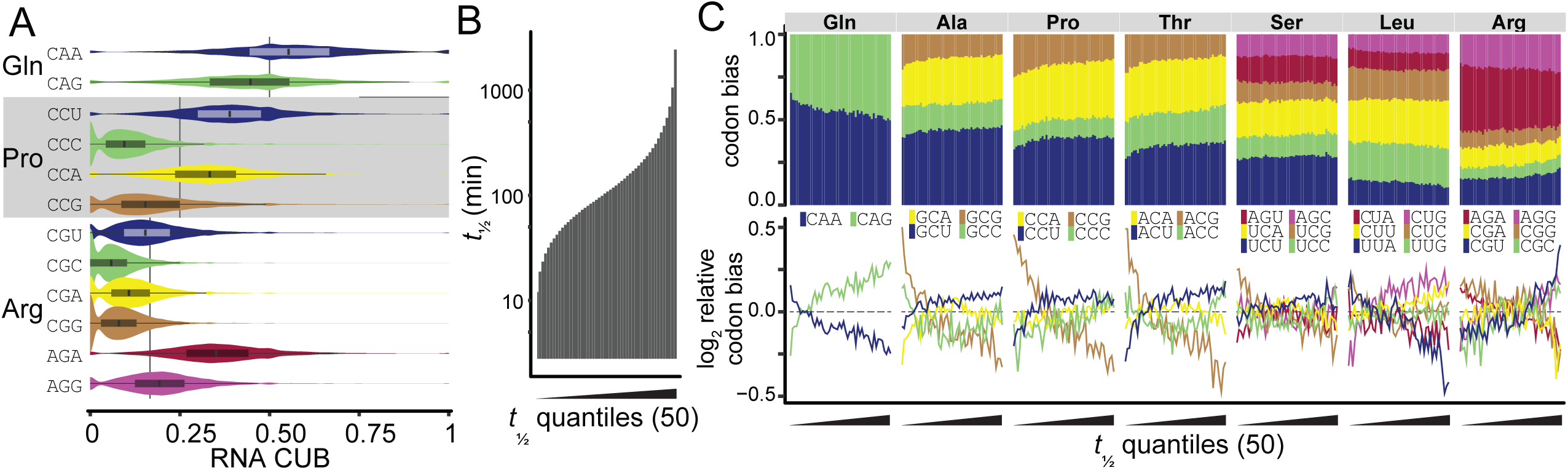
Codon usage differs for mRNAs with short and long *t*_1/2_s. (A) Distribution of Gln, Pro, and Arg CUB of 16,749 individual transcripts expressed in 5-d-old Arabidopsis thaliana seedlings. CUB was calculated for each transcript; if a transcript did not encode a given amino acid it was not counted in the distribution. Boxplots were overlaid on violin plots to show codon bias distributions. Vertical black lines behind violin plots represents the expectation if codon usage was random. (B) Median mRNA *t*_1/2_ of each of 50 quantiles of 16,749 transcripts from whole 5-d-old Arabidopsis seedlings [4]. (C) Codon bias (top) and log2 quantile codon bias relative to genome-wide codon bias (bottom) of 50 mRNA *t*_1/2_ quantiles (increasing left to right as in A). Codons of Gln, Ala, Pro, Thr, Ser, Leu, Arg and show trends across *t*_1/2_ quantiles.

To determine whether differences in CUB across mRNAs were related to stability, we organized mRNAs by increasing *t*_1/2_ and separated them into 50 quantiles (or bins) (Figure 1B), and for each bin, we calculated mean mRNA CUB. Across all bins, CUB varied by as much as 16% (Gln codons CAA and CAG), or as little as 2% (Gly GGC) (Supplemental Figure 1B). To visualize trends, codon use in each bin was normalized to genome-wide usage (log_2_ mean bin codon usage/mean codon usage of all genes) (Figure 1C, Supplemental Figure 1B). This data transformation revealed that for some amino acids (*e.g.*, Gln, Ala, Pro, Thr, Ser, Leu, Arg) a distinct relationship was apparent, but for others (*e.g.*, Glu, His, Gly) there was little to no relationship between *t*_1/2_ and CUB. Using linear regression, we found 46 codons that had a significant relationship with decay rate (*p* < 0.01), and some codons explained up to 3% of the variance (Supplemental Data 1). Of note, rare codons were no more likely to be associated with *t*_1/2_ than common codons (Figure 1C). For example, CCC (Pro) usage was rare (median of 9.5%) but mRNAs with higher usage of CCC (relative to the whole transcriptome) had longer *t*_1/2_s, whereas AGA (Arg) was the most common codon (median of 35%), but mRNAs with higher AGA usage (relative to the whole transcriptome) tended to have shorter *t*_1/2_s. Thus, high codon usage per se was not associated with mRNA stabilization and vice versa.

### Codons can be used to roughly predict mRNA decay rate

The association between CUB and mRNA stability suggested that rates of synonymous codon occurrence within an ORF might be predictive of its mRNA decay rate. To explore this possibility, we assessed whether codon composition of ORFs was linked with mRNA stability and reflected the trends in CUB. To test this, we calculated each codon’s impact as a codon stabilization coefficient (CSC [24]), defined here as the Pearson correlation between transcript codon frequency and -log10 decay rate. Frequency was positively correlated with stability for some codons, while for others it was negatively correlated (Supplemental Figure 2). For example, CUG (Leu) had higher frequency in stable transcripts (positive CSC), while UUA (Leu) was used more in unstable mRNAs (negative CSC) (Figure 2A). Across all 64 codons, we observed correlations ranging from -0.18 for UCG (Ser) and AGA (Arg) to the strongest correlation of 0.23 for GCU (Ala) which alone explains 4.84% of decay rate variance. To determine whether CSCs were dependent on nucleotide sequence or nucleotide composition, we performed a permutation test by scrambling ORF sequences and used the resultant triplet nucleotide frequencies to compute stabilization coefficients (SCs) similar to those of codons (Figure 2B, Supplemental Data 2). This test showed that 53 CSCs were significant (p < 0.05, outside of the 95%CI of the permutations’ SCs) suggesting it is highly unlikely that CSC values are random or due to nucleotide composition (Figure 2A). Instead, codons are likely to influence mRNA stability.

**Figure 2.**
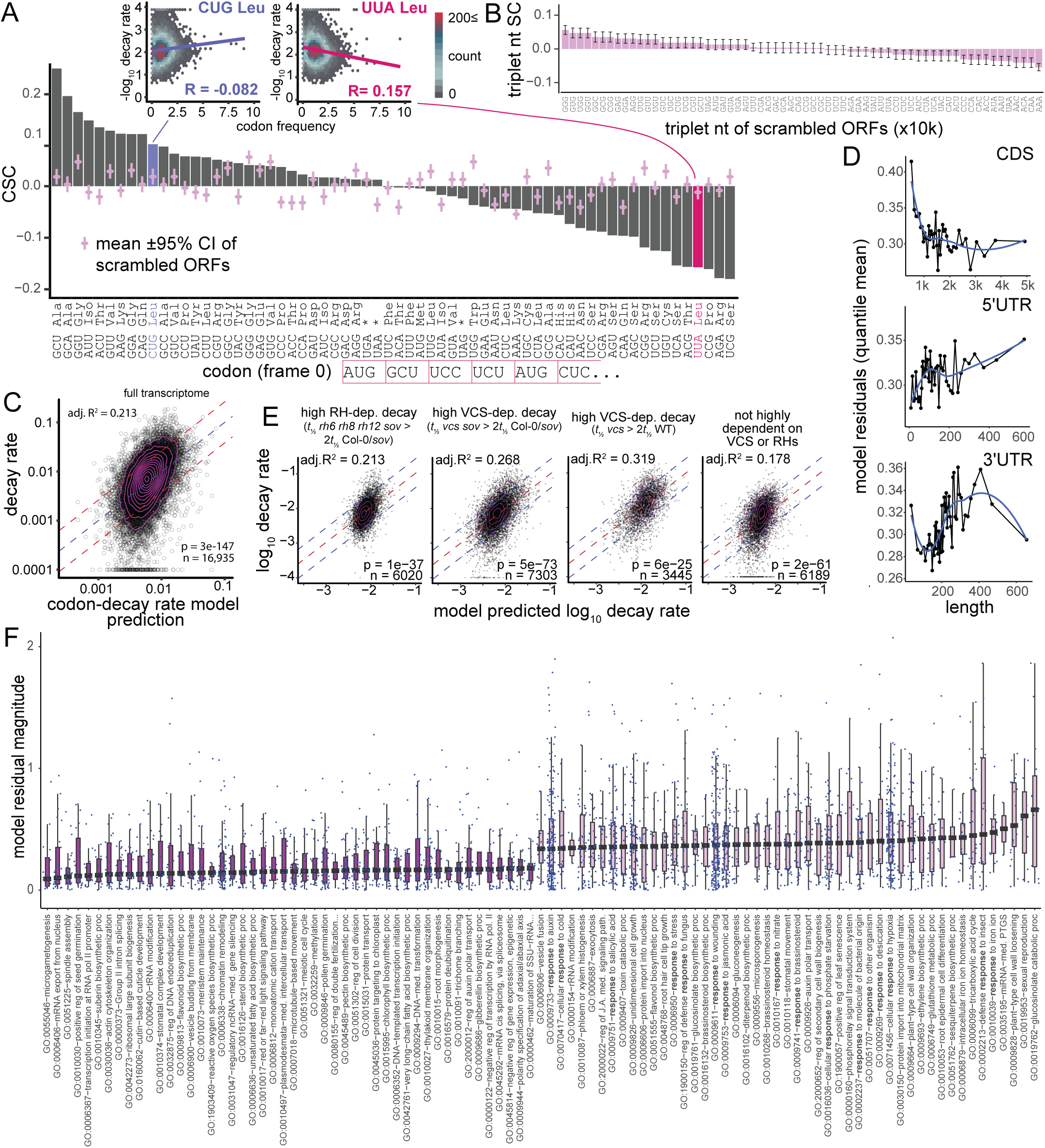
Codon frequencies can roughly predict mRNA decay. (A) Codon stabilization coefficient (CSC, Pearson correlation of codon frequency and -log10 decay rate for all nuclear-encoded and protein-encoding RNAs). CSCs are compared with CSCs of the scrambled ORF triplet nucleotides ±95% CI (purple). CSC values of UUA (Leu) and CUG (Leu) are colored and labeled for comparison with insets. Insets: scatter-density plot of codon frequency (as percent) with log10 decay rate for two codons encoding Leu; CUG has a negative correlation (blue regression line) indicating that it is used more in stable mRNAs, and UUA has a positive correlation (pink regression line) indicating less use in stable mRNAs. Points are colored by mRNA density as indicated. (B) SCs of triplet nucleotides from scrambled transcript ORF sequences. Each ORF was scrambled to calculate triplet nucleotide frequencies, then based on all scrambled mRNAs stabilization coefficients (SCs) were calculated as was done for codons. Mean±95% confidence interval (CI) was estimated from SCs calculated from 10,000 permutations of ORF scrambling. (C) Multiple regression model performance. Scatter plot comparing the codon frequency model-predicted decay rate (log10) with the measured decay rate (log10). Magenta contours indicate zones of equal point density. Blue dotted line indicates the line of perfect prediction; red dotted lines are ±1 SD of the residuals. (D) Composite codon model residuals were averaged for each of 50 bins of RNAs for CDS length, 5’ UTR length, and 3’ UTR length. Lower residuals indicate more accurate decay rate prediction based on codon frequencies. (E) Composite codon model fit to subsets of RNAs based on RH- or VCS-dependence. (F) Biological Process GO categories (>20 gene members) with the largest (pink) and smallest (blue) codon-decay model residuals. Boxplots are ordered by median residual.

We next sought to understand how much combined codon composition of ORFs determine decay rates. To determine how much of the total variance in transcriptome decay rates might be explained by codons, we modeled mRNA decay rate as a function of all codon frequencies in each transcript. Following backward stepwise regression (p < 0.01), the final linear model used codon frequencies of 39 of the 61 amino-acid-encoding codons; of these, 19 coefficients were positive, associated with faster decay, and 20 were negative, associated with slower decay (Supplemental Figure 3A, Supplemental Data 3). Larger coefficients, positive or negative, indicated a stronger influence on decay rate. Hereafter, we refer to this model as the codon-decay rate model. Model performance was visualized as a scatter plot of predicted decay rate values versus actual measured values (Figure 2C). Overall, codon frequencies explained 21.3% (adj. R^2^ = 0.213, p = 3x10^-147^) of transcriptome-wide mRNA decay rate variance. Thus, codon content was a strong predictor of mRNA decay rates, consistent with COMD as one of several mechanisms governing mRNA stability in Arabidopsis [16].

### Characteristics of mRNAs with strong codon-decay associations

Performance of the codon-decay rate model varied by mRNA, suggesting that COMD might be the main decay rate determinant for some RNAs, but not others. To identify features that distinguished mRNAs for which codons were good predictors of decay rate, we assessed the relationship between model accuracy and mRNA physical features – specifically, coding sequence (CDS) length, 5’-UTR length, and 3’-UTR length. We divided mRNAs into 50 bins of length for each feature and used the model prediction error (residual) to indicate the accuracy of the codon-predicted rate. Comparing median residual size across the 50 bins, we found that the model had mildly better predictions (smaller residuals) for mRNAs with CDS > ∼750 nt, short 5’-UTRs (<35 nt), and moderately short 3’-UTRs (100-200 nt) compared with those with different lengths (Figure 2D). Modeling only mRNAs with CDS > 750nt (86%) increased explained variance to 22.4%. These data suggest that codon composition is a better predictor of decay rate for mRNAs with long CDS and short UTRs.

To identify functional categories of mRNA for which codon usage was a good predictor of decay, we assessed gene ontologies (GOs). We calculated the median codon-decay rate model residual for all GOs with more than 20 members (790 GOs). GOs with the most predictive codons (smallest residuals, 6%, 48 GOs) were mostly related to development and morphogenesis (19 GOs) or gene expression and regulation (17 GOs), while none were characterized as responsive RNAs (Figure 2F). By contrast, many of the GOs with the least predictive codons (largest residuals, 6%, 49 GOs) were related to plant responses to environment and stress (13 GOs) or plant hormone response and metabolism (9 GOs). Thus, decay rates of mRNAs that are involved in the fundamental molecular machinery of gene expression and growth are more likely to be determined by codon content and mRNAs characterized as responsive to hormones, environmental interactions, or stress responses are predominantly determined by mechanisms unrelated to codon content.

We also asked whether specific decay pathways were involved in codon-associated decay rates. We expected that COMD would initiate during translation and, therefore, require decapping to allow XRN4-mediated co-translational degradation [15,39–43]. Similarly, COMD might also require Dhh1-like genes in Arabidopsis, *RNA helicase* (*RH*) *6*, *RH8*, and *RH12*, given the role of yeast Dhh1 in decapping recruitment [44–47]. To test these ideas, we evaluated the relationship between combined codon composition and decay rate for mRNAs that have high dependence on the decapping complex scaffold VCS, high dependence on Dhh1-like RHs, and mRNAs that degraded by other pathways. High dependence was defined as a 2-fold decrease in in decay rate in mutants (*vcs-7* or *rh6812* triple mutant) compared with control plants [4,48]. For mRNAs with high VCS dependence, codons explained decay variance better than those of the whole transcriptome – 27% in the Col-0 (*sov*) background and 32% in the background with a functional SOV (3’-to-5’ exoribonuclease) (Supplemental Data 4, 5), whereas mRNAs with high RH-dependent decay were similar to the whole transcriptome (21% variance explained) (Figure 2E, Supplemental Data 6). By contrast, codons of mRNAs with decay not dependent on VCS or RH only explained 17% of decay rate variance (Supplemental Data 7). These data are consistent with a specialized role for mRNA decapping in codon-mediated decay, but not Dhh1-like RHs or other decay pathways in Arabidopsis.

### Manipulation of synonymous codon usage confers predictable changes to mRNA decay rate

We tested the codon-decay rate model predictions by generating two synonymous alleles of YFP. Codons used more in stable or more in unstable transcripts were used to generate *YFP^stCod^* and *YFP^unCod^*, respectively. These alleles were expressed under control of the Arabidopsis *UBIQUITIN 10* (*UBQ10*) promoter (*proUBQ10* [49]) (Figure 3A) in stably transformed Arabidopsis plants to assess the impact of recoding on stability. Because differences in mRNA decay rate might be reflected in protein and mRNA accumulation, we compared recoded YFP protein and mRNA abundances. Root tip fluorescence was significantly higher in *YFP^stCod^* plants than *YFP^unCod^* plants (Figure 3B and 3C). mRNA abundance in seedlings was >10-fold higher in *YFP^stCod^* than *YFP^unCod^*(Figure 3D). Because both alleles used the same *UBQ10* promoter, their differing mRNA abundances likely resulted from differing decay rates. To test this, we measured mRNA *t*_1/2_ using a transcription termination assay. The mRNA *t*_1/2_ of *YFP^stCod^*was 64 min whereas that of *YFP^unCod^* was 12 min (Figure 3E). This 5-fold decay rate difference was specific to the transgenes as native gene transcripts *ACT2* (*t*_½_ 19.25 h) and *ZAT10* (*t*_½_ 13 min) decayed at rates similar to previous measurements in both transgenic genotypes [4] (Supplemental Figure 3B). These results were consistent with codon usage affecting protein levels, and mRNA abundance and stability.

**Figure 3.**
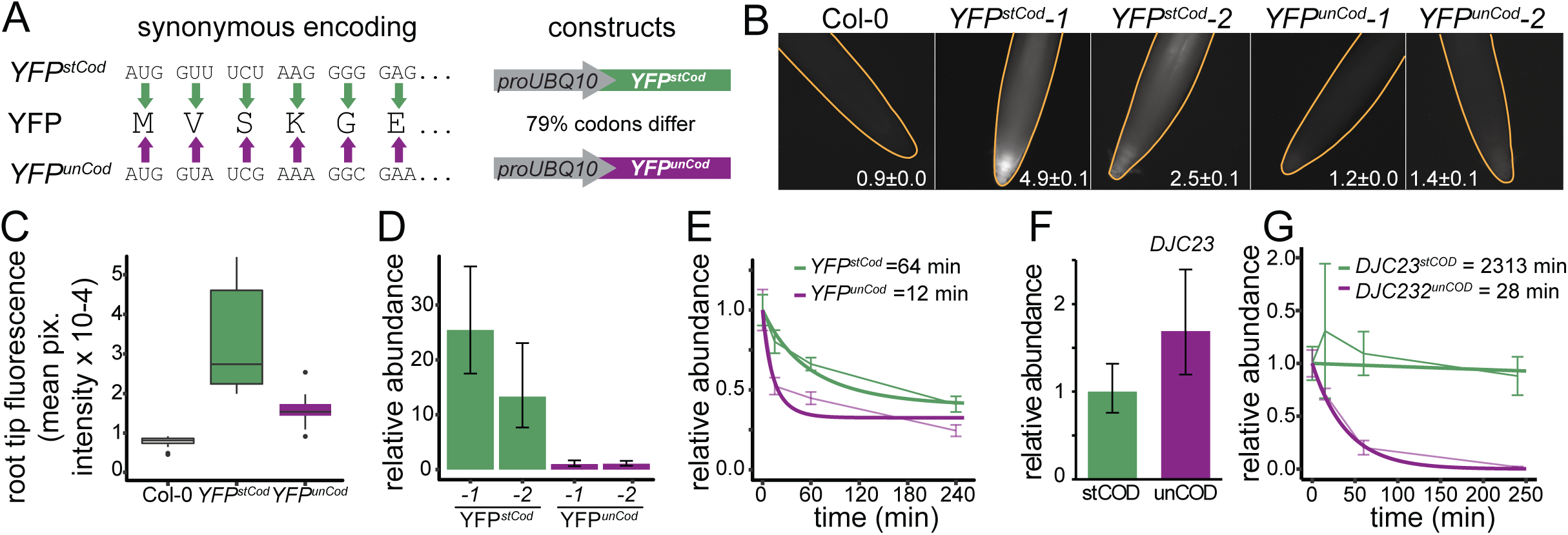
Model-directed synonymous encoding predictably alters mRNA stability. (A) Diagram of synonymous encoding of YFP for maximum predicted stability using codons associated with stable mRNAs (stCod) or minimal predicted stability using codons associated with unstable mRNAs (unCod). Each were fused to the promoter of *UBQ10*. (B) Representative fluorescence images of root tips (outlined in orange) for an untransformed plant (Col-0) and two independent insertion lines each of *YFPstCod* and *YFPunCod*. White numbers are mean pixel intensity x 10-4 ±SE (n = 4 plants). (C) Boxplot of root tip fluorescence of untransformed plants (n = 16), YFPstCod (n = 5 independent lines, 2 plants each), and YFPunCod (n = 11 independent lines, 2 plants each). (D) Relative *YFP* mRNA abundance measured by qRT-PCR normalized to *YFPunCod*-1 (mean±SE, using both *ACT2* and *eEF1Bγ* reference mRNAs). (E) *YFP* mRNA abundance after treatment with cordycepin to block RNA synthesis (thin lines; mean±SE, n = 4 biological replicates each of 2 independent insertion lines). mRNA abundance was normalized to stable reference mRNAs: eEF1Bγ, UBC10, and VHA-A and to T0 samples. mRNA decay (α) and decay of decay (β) rates were estimated by maximum likelihood modeling (thick lines). Half-lives (*t*_1/2_s) were calculated from these estimates and are indicated in the key. (F, G) Relative mRNA abundance of *XVE >> DJC23stCOD* and *XVE >> DJC23unCOD* mRNAs were measured after 24 h induction by beta-estradiol and normalized to reference gene *UBC10* (F). Relative decrease in abundance after cordycepin treatment (G, thin lines; mean±SE, n = 4 biological replicates each of 2 independent insertion lines) and modeling of mRNA decay (α) and decay of decay (β) rates were estimated by maximum likelihood modeling (thick lines). *t*_1/2_s were calculated from these estimates and are indicated in the key.

We also tested the codon-decay rate model by recoding *DJC23,* an endogenous gene with a role in protein folding. We changed synonymous codons to make two versions, *DJC23^stCod^* and *DJC23^unCod^*, and placed these under control of the estradiol-inducible *XVE* promoter system [50,51]. We induced expression for 24 h, obtained similar RNA abundances (Figure 3F) and measured decay rates. We found an 83-fold difference in *t*_1/2_ - 2,313 min for *DJC23^stCod^* and 28 min for *DJC23^unCod^* (Figure 3G). Together, these results indicate that selected codons had a large impact on RNA stability and that codon frequency correlations with mRNA decay rates are predictive of these impacts in plant genes.

### Codon-independent features also affect mRNA decay rate

Our data is consistent with the COMD model in Arabidopsis thus far. To attempt to disprove the model, we tested these two additional predictions. In yeast and humans, COMD occurs during translation. A low-optimality codon is decoded more slowly in the ribosome A-site, allowing tRNA to vacate the E-site. The ribosome with empty E- and A-sites changes conformation, and the N-terminal domain (NTD) of Not5 in budding yeast (*Saccharomyces cerevisiae*) or CNOT3 in humans binds the empty E-site in this conformation [42,52]. In human cells, specific Arg tRNAs in the P-site can also directly recruit NTD interaction [52]. Not5/CNOT3 then recruits the CCR4-NOT complex for deadenylation [42]. This model leads to the predictions that in plants (1) the Not5/CNOT3 NTD is conserved, and (2) our observed sequence (codon) correlations with decay must depend on the translation reading frame.

To assess conservation of the NTD, we compared amino acid sequences of the Arabidopsis ortholog [53] with yeast Not5 and human CNOT3. Not5 and AtNOT3 NTDs shared 51% identity and 69% similarity (Supplemental Figure 4A, 4B[54]). Twelve of 15 Not5-NTD residues that mediate ribosome interaction were also identical or similar [42]. Conservation of these residues in 9 additional diverse eukaryotes suggests their importance. Additionally, 4 of 5 residues that mediate P-site Arg-tRNAs interactions with human CNOT3 NTD were identical in Arabidopsis [52]. Further evidence of conservation of this interaction is found in Arg tRNA sequences. Five nucleotides in these tRNAs mediate interaction with CNOT3 NTD. All five were identical in Arabidopsis Arg tRNAs with UCU or CCG anticodons (8 unique tRNA sequences) but not any other Arabidopsis tRNAs including those of the 13 other Arg tRNAs (188 unique sequences) (Supplemental Data 8). Moreover, the codons decoded by the UCU or CCG tRNAs (AGA and CGG) had large negative CSC values suggesting they were used in unstable mRNAs (Figure 2A), and, as in human cells, Arabidopsis mRNAs with high use of these codons were enriched in *mitochondrial large ribosomal subunit* (p-adj. 0.014) (Supplemental Data 9). Together, these observations strongly suggested that the mechanism for sensing codon optimality and specific Arg-tRNAs in the P-site is conserved in plants and has an ancient origin.

To determine whether codon-mRNA decay rate correlations were determined through translation of codons and not via other sequence features, we assessed the dependence of these correlations on the translation reading frame. If translation is critical for decay, then the correlations between frequencies of frameshifted codons and mRNA decay rates for the +1 and +2 frames should be different than those of codons, and more similar to those calculated for scrambled mRNAs (Figure 2B). We tested this prediction. SCs of the +1 and +2 reading frames were calculated in the same way as codons (CSCs), namely using Pearson correlation of out-of-frame (+1 and +2 frames) triplet nucleotide frequency and -log10 mRNA decay rates. Surprisingly, out-of-frame triplet nucleotide frequencies were correlated with mRNA decay as strongly as codons, and differed from those of the scrambled mRNA sequences (56 significant triplets in each frame, p < 0.05, permutation test) (Figure 4A). Moreover, five out-of-frame triplet nucleotides had even stronger correlations than any codon. This lack of reading frame dependence in sequence-decay correlations conflicts with the predictions of the COMD model and suggested that codons themselves might not be the only mRNA sequence influence on RNA *t*_1/2_s.

**Figure 4.**
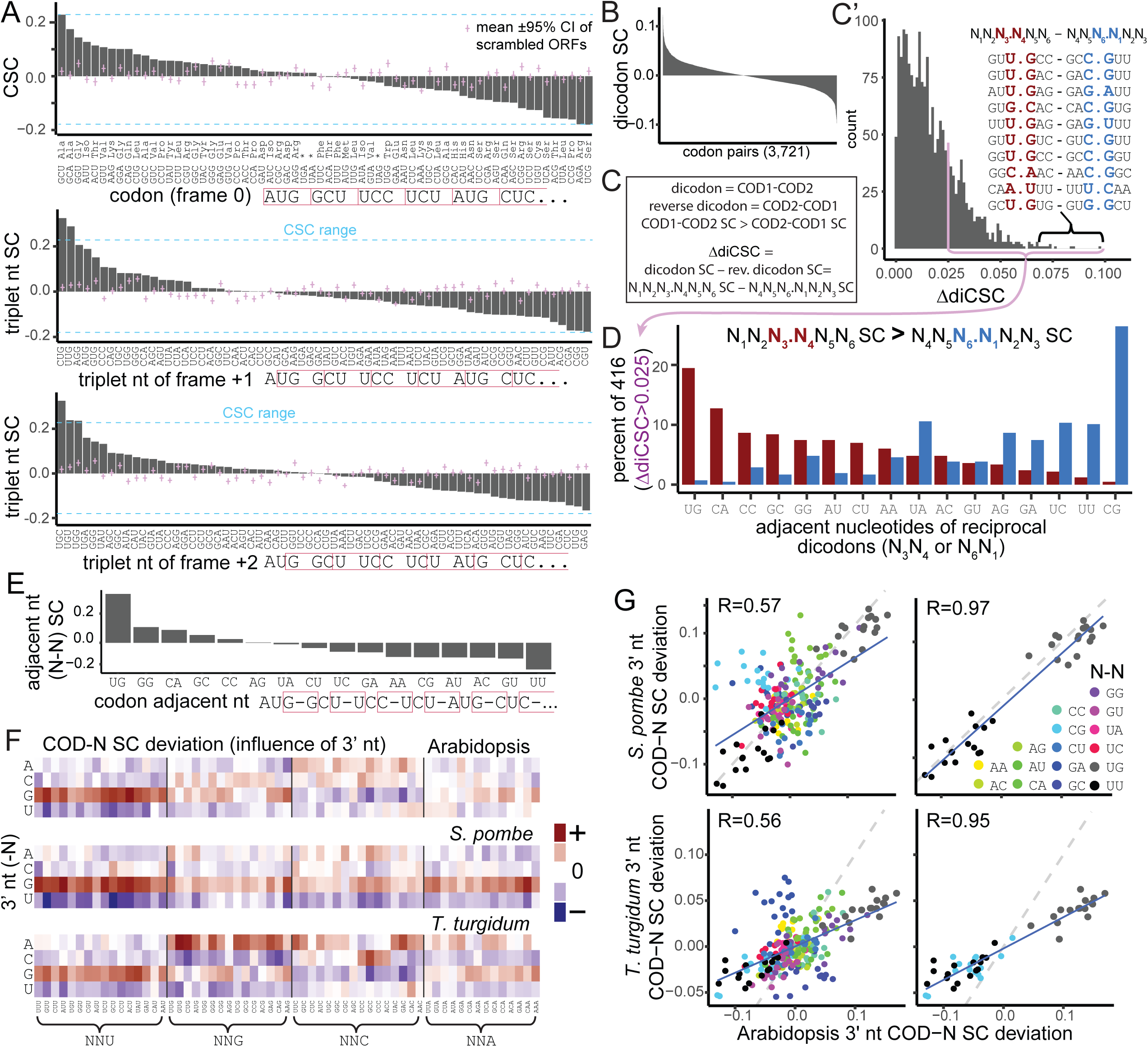
NCANs are strongly associated with decay rate independently of codons. (A) Triplet nt SCs for frame-shifted ORFs (frame +1 and +2). Triplets are ordered in each plot by SC. SCs are compared with those of the scrambled ORF triplet nucleotides ±95%CI. (B) Dicodon stabilization coefficient (SC; dicodon frequency ∼ log10 WT decay rate Pearson correlation) bar plot for 3,721 dicodons (excluding stop codons). Dicodon SCs range -0.123 to 0.0971. (C) Definition of dicodon, reverse dicodons, a dicodon nucleotide index, and the reciprocal dicodon SC difference (ΔdiCSC). The reverse order was defined by the smaller diCSC and subtracted from the larger diCSC. (C’) Histogram of SC difference for reciprocal dicodon (the same two codons but in reverse order). The order of the pair was selected such that COD1.COD2 SC was greater than COD2.COD1 so that the SC difference would be positive. The dark magenta line at 0.025 indicates a shift of 11.3% of the total dicodon SC range and separates 22.7% of 1830 reciprocal dicodons. The 10 dicodons with the highest SC difference are listed. (D) Adjacent nucleotides of are associated with the SC difference of reciprocal dicodons The adjacent nucleotides from 416 reciprocal dicodons with a ΔdiCSC > 0.025 were assessed. The fraction of all adjacent nucleotides from the dicodon orientation with the larger SC (red) are compared with the same from the reverse orientation and lower dicodon SC. (E) NCANs (N-N) stabilization coefficients. (F) The effect of the 3’-neighboring nucleotides on codon SCs. Heatmap of COD-N SC deviation (mean COD-N SC by codon - COD-N SC) for Arabidopsis, *T. turgidum*, and *S. pombe*. (G) Scatter plot comparison of Arabidopsis and *S. pombe* 3’-adjacent nucleotide influences on COD-N SCs (mean COD-N SC by codon - COD-N SC).

### Neighboring codons’ adjacent nucleotides also correlate with mRNA decay rate

We considered that other sequence motifs or features of codon sequence might influence ribosome decoding and therefore decay rate. For example, codons context can influence translation rate in both eukaryotes and prokaryotes [55–57] and using successive codon pairs (dicodons) composition improved predictive indices for mRNA expression compared with codon composition alone [58]. To determine whether the SCs of the +1 and +2 reading frames resulted from non-additive effects of neighboring codons, we calculated dicodon SCs (diCSC, i.e., dicodon frequency ∼ -log10 decay rate Pearson correlation) (Figure 4B, Supplemental Figure 5A, Supplemental Data 10) to compare them with their component codons CSCs.

We assessed the influence of codon context on diCSC by evaluating whether the order of codons had any effect on diCSC magnitude. We reasoned that if codon influence is merely additive, then diCSCs should represent the average influence of the two component codon, regardless of the codon order. diCSCs of reciprocally ordered dicodons were compared by calculating their difference (ΔdiCSC, 1,830 excluding stop codons) (Figure 4C, Supplemental Data 11). Consistent with little influence of codon context, most reciprocal dicodons had small differences; however, some were large (>0.025, cutoff based on the inflection of the distribution, Figure 4C’). When examining the ten dicodons with the largest ΔdiCSCs, we noticed that the adjacent nucleotides (N_3_N_4_) of the dicodons (COD1.COD2, larger diCSC) appeared over-represented for UG (7/10, p = 2x10^-10^, hypergeometric test). By contrast, in the adjacent nucleotides (N_6_N_1_) of the reverse dicodons (COD2.COD1, smaller diCSC), CG was over-represented (6/10, p = 3x10^-8^, hypergeometric test). This suggested that dicodon adjacent nucleotides might explain non-additive codon effects of diCSCs. We expanded this analysis to include reciprocal dicodons with large ΔdiCSCs (> 0.025, 22.7%) and found a striking trend; UG were the adjacent nucleotides in 19.5% of dicodons with the larger diCSC codon order (N_3_N_4_) and 0.7% of the reversed codon order (smaller diCSCs, N_6_N_1_) (Figure 4D). The reverse of this pattern was observed for adjacent nucleotides CG; 26.4% of dicodons with the reverse order had CG (N_6_N_1_), whereas CG was present in only 0.5% of dicodons with larger diCSC codon order (N_3_N_4_). Other adjacent nucleotides also showed tendencies to occur in the larger or smaller diCSC codon orders (e.g., UA, UC, UU as N_3_N_4_ and CA as N_6_N_1_), suggesting that neighboring codons’ adjacent nucleotides (NCANS) might impact mRNA stability.

To determine whether specific adjacent nucleotides were associated with stable or unstable transcripts, we calculated a SC based only on NCANs (frequency of dicodon’s adjacent nucleotides ∼ -log10 decay rate Pearson correlation) (Figure 4E, Supplemental Data 12). Adjacent nucleotide SCs ranged from -0.099 (C-G) to 0.335 (U-G) suggesting a range of association with decay rate. A multiple regression model applied to adjacent nucleotide frequency (as was to codon frequency) revealed that NCANs alone explained 16.7% of decay rate variance (p < 2.2e-16). Because codon order determines adjacent nucleotide pairs, codon context appears to be an additional factor in RNA stability.

Given that NCANs alone and codons alone both show strong correlations with decay rate, we tested the impact of the codons together with NCANs by calculating the SC of each codon plus one neighboring nucleotide either 5’ or 3’ (which we referred to as N-COD or COD-N, respectively) (Supplemental Data 13). We calculated the effect of the additional nucleotides by subtracting the N-COD or COD-N SC from the mean of each codon (i.e., [A, U, C, and G]-COD or COD-[A, U, C, and G] SCs) and visualized these values as a heatmap. Depending on what adjacent nucleotide pairs were formed, the additional nucleotides had consistent positive or negative influence. For example, UG always had a strong positive effect on the SC relative to other adjacent nucleotide pairs, and CG always had a negative effect, and similar but weaker effects were observed for other adjacent nucleotide pairs. Applying a multiple regression model of log10 decay rate to COD-N frequencies instead of just codon frequencies resulted in an increase in explained decay rate variance to 26.4% for COD-N frequencies compared with 21.3% for that of codons alone (Supplemental Data 14). This boost in model performance supports that NCANs as an additional factor influencing RNA stability.

### NCANs are associated with RNA stability in distant eukaryotes

To determine whether a relationship between NCANs and RNA decay is found in eukaryotes we evaluated published datasets from multiple species. As a base line for comparison, we first compared codon optimality between distant eukaryotic organisms using the CSC metric. Arabidopsis CSCs were positively correlated to varying degrees with budding yeast (*Saccharomyces cerevisiae*, R = 0.34) [24], fission yeast (*Schizosaccharomyces pombe*, R = 0.55) [59], zebrafish (*Danio rerio,* R = 0.61) [26], frog (*Xenopus tropicalis*, R = 0.49) [26], and wheat (*Triticum turigidum*, R = 0.51) [3], but not human (R = -0.20) [31] suggesting some similarity in codon optimality across eukaryotes (Supplemental Figure 5C).

We next calculated COD-N SCs for each of these species. In *S. cerevisiae*, there was no specific pattern associated with adjacent nucleotide pairs (Supplemental Figure 5D - heatmap of COD-N), but in *S. pombe* and *T. turgidum*, we found adjacent nucleotide influences on SCs similar to those in Arabidopsis (R=0.57 and R=0.56 respectively, Figure 4F, 3G). Specifically, the U-G and U-U adjacent nucleotides had similar strong effects in fission yeast (R=0.97), whereas, C-G and A-G had large differences. In wheat, U-G, C-G, and U-U SCs were very similar (R=0.95) while G-A had a uniform but different effect compared with Arabidopsis. There was clear influence from the adjacent nucleotides on the codons, despite these differences. Applying the codon-decay model to wheat revealed that codons explained 13.1% (p-value = 5x10^-9^) decay variance, and including codons’ 3’-adjacent nucleotides in the COD-N-decay model increased the explained variance to 16.8% (p-value = 2x10^-57^) (Supplemental Data 15-16). We also calculated COD-N SC data for frog and zebrafish (Supplemental Figure 5D). Overall, 3’-adjacent nucleotide influences weakly correlated with those of Arabidopsis (R = 0.258 and R = 0.365, respectively) (Supplemental Figure 5E), but again, U-G and U-U consistently had similar influences on COD-N SCs of these two species (R of 0.81 and 0.69, respectively), suggesting some conservation of influence by NCANs on decay in distantly related eukaryotes.

## Discussion

We provide evidence that codon-optimality-mediated decay functions in Arabidopsis and that additional sequence features appear to influence the codon-decay relationship. Codon content explained 20-30% of decay rate variance, depending on the mRNA population. This finding along with conservation of the AtNOT3 codon optimality sensor domain and a causal relationship between codons and decay rates suggests that mRNA decay can and frequently does initiate based on codon optimality in Arabidopsis. Moreover, we discovered strong correlative data that NCANs might also influence mRNA decay rates. This phenomenon was clearly observed in Arabidopsis, wheat and fission yeast, suggesting that codon context could influence mRNA decay rate in distant eukaryotes. These findings could be useful in future studies for design of coding sequences of synthetic constructs with specific RNA stabilities in Arabidopsis.

### Codon-optimality-mediated decay is one of multiple pathways that determine decay rates

Depending on the mRNA population codon frequencies accounted for 20-30% of decay rate variance in Arabidopsis (Figure 2C and Figure 2E). While this was a substantial proportion of variance, this means other mechanisms also substantially contribute to RNA decay rate determination in Arabidopsis. Wu et al. [3] suggested that, in tetraploid wheat, 3’-UTR structural motifs played a major role in mRNA stability, with some motifs having a stabilizing effect and others a destabilizing effect. Although codons explain more decay rate variance in Arabidopsis than in wheat, 3’-UTR motifs likely also function in Arabidopsis. The Arabidopsis codon-decay model had better decay rate prediction for mRNAs with longer CDS, a higher decay dependence on decapping, and moderately short 3’ UTRs (Figure 2D). With longer CDS, there are more codons and a higher chance that one of the codons will recruit NOT3. NOT3 recruits deadenylation which is often followed by decapping and can allow co-translational decay. This was consistent with decapping dependent mRNAs having a stronger association between codons and decay (Figure 2E). Longer 3’ UTRs have more likelihood of containing mRNA stability motifs that might influence decay rate independently of codon usage. For example, cis-elements, RNA G-quadruplex structures, and miRNA target sites can all affect RNA stability, and they commonly occur in 3’-UTRs [3,60,61]. Although we removed experimentally determined miRNA targets from our model, there might be many more miRNA targets as approximately half of all genes are predicted to be miRNA targets [60]. Nonsense-mediated decay (NMD) also affects decay rates of many mRNAs. Drechsel et al. [62] suggested that up to 17% of protein coding genes have splice variants that are targeted by NMD - especially for mRNAs with long 3’-UTRs providing another possible explanation for increasing codon model residuals with longer 3’UTRs.

Because plants are sessile, one of the main ways they respond to myriad stimuli is through tailored genetic programs that change RNA abundances. Responsive RNA abundance is often controlled through dynamic transcriptional regulation, but dynamic RNA decay regulation can also contribute to changes in RNA abundance. It is conceivable that RNA decay rates could be regulated through changes in codon optimality. For example, changes in tRNA expression can drive changes in gene expression and cancer progression [63] and decoding rates of specific codon can change in response to a stimulus [64]. Alternatively, optimality of specific codons might also be regulated through shifts in tRNA gene expression (i.e., from one tRNA gene to another with different sequence or folding structure but the same anticodon, Supplemental Data 8) or changes in aminoacyl tRNA synthetase activity. However, these modes of regulation would influence cohorts of RNAs. COMD seems a better fit for RNAs with more consistent abundances because we found that codon content does not predict decay rate as well for mRNAs with abundances that are responsive (Figure 2F). However, more studies are needed to understand the effects of specific tRNA modifications as well as levels and dynamic changes in tRNA charging; our current understanding of these variables is limited [16,65,66].

### How might adjacent nucleotides influence mRNA stability?

Codon stabilization coefficients were good predictors of codon influence on mRNA stability (Figure 3), thus it is reasonable to expect that the stabilization coefficients of NCANs also indicate influence on Not5/NOT3 recruitment. Understanding the dynamics of Not5/NOT3 recruitment due to codon optimality gives us some hint. Low codon optimality in the ribosome A-site influences the probability of Not5 NTD recruitment to the E-site because it slows decoding-time and increases the probability of the empty E- and A-site ribosome conformation required for binding [42]. However, Not5/NOT3 recruitment to ribosomes appears to be a rare event relative to continuous translation. In other words, a codon with low optimality only results in decay for a very small percentage of its decoding events. Each coding sequence is composed of many codons with varying levels of optimality, but only one codon is required to recruit Not5/NOT3 and initiate decay of the mRNA molecule. For a rough approximation of the frequency of Not5/NOT3-mediated decay initiation, we can assume that for a 600-codon ORF being actively translated at a modest average rate of 5 codons s^-1^, one ribosome would take 2 min to produce one protein [67]. In a pool of 100 mRNA copies being actively translated in polyribosomes with an average ribosome spacing of 90 nt, 2,000 proteins could be produced in 2 min [68]. For a common plant *t*_1/2_ of 139 min, only one of these 100 mRNA copies would need to be targeted for degradation during the 2 min round of translation, i.e., only 1 of 1,200,000 translated codons (2,000 full ORF translations) would need to recruit Not5/NOT3. For a short 4 min *t*_1/2_, still only 1 of 34,286 translated codons (57.1 full ORF translations) would trigger degradation.

Given this low frequency estimate of Not5/NOT3 recruitment, it is plausible that subtle influences by variables other than charged tRNAs concentrations could have a large effect on the rate of Not5/NOT3 recruitment at any of a number of positions in the ribosome’s codon register. One of these variables has already been demonstrated by Zhu et al. [52]; some Arginine tRNAs in the ribosome P-site can directly enhance Not5/NOT3 affinity and therefore its frequency of recruitment. Adjacent nucleotide of neighboring codons might subtly shift affinities of charged tRNAs in the A-site, uncharged tRNAs in the E-site, the speed of ribosome conformational change, or Not5/NOT3 binding affinity (Figure 5C). For example, is it possible that nucleotide three of the codon in the E-site influences the rate of uncharged tRNA release in a manner dependent on the next 3’-nucleotide in the P-site, or in the same way affects the recruitment of Not5/NOT3 interaction? Might nucleotide three of the P-site codon affect charged tRNA recruitment in the A-site depending on the adjacent 3’-nucleotide? More studies are needed to determine what might account for the adjacent nucleotide-decay correlation of codon sequences and to disentangle the effects of codons alone from the effect of their context.

**Figure 5.**
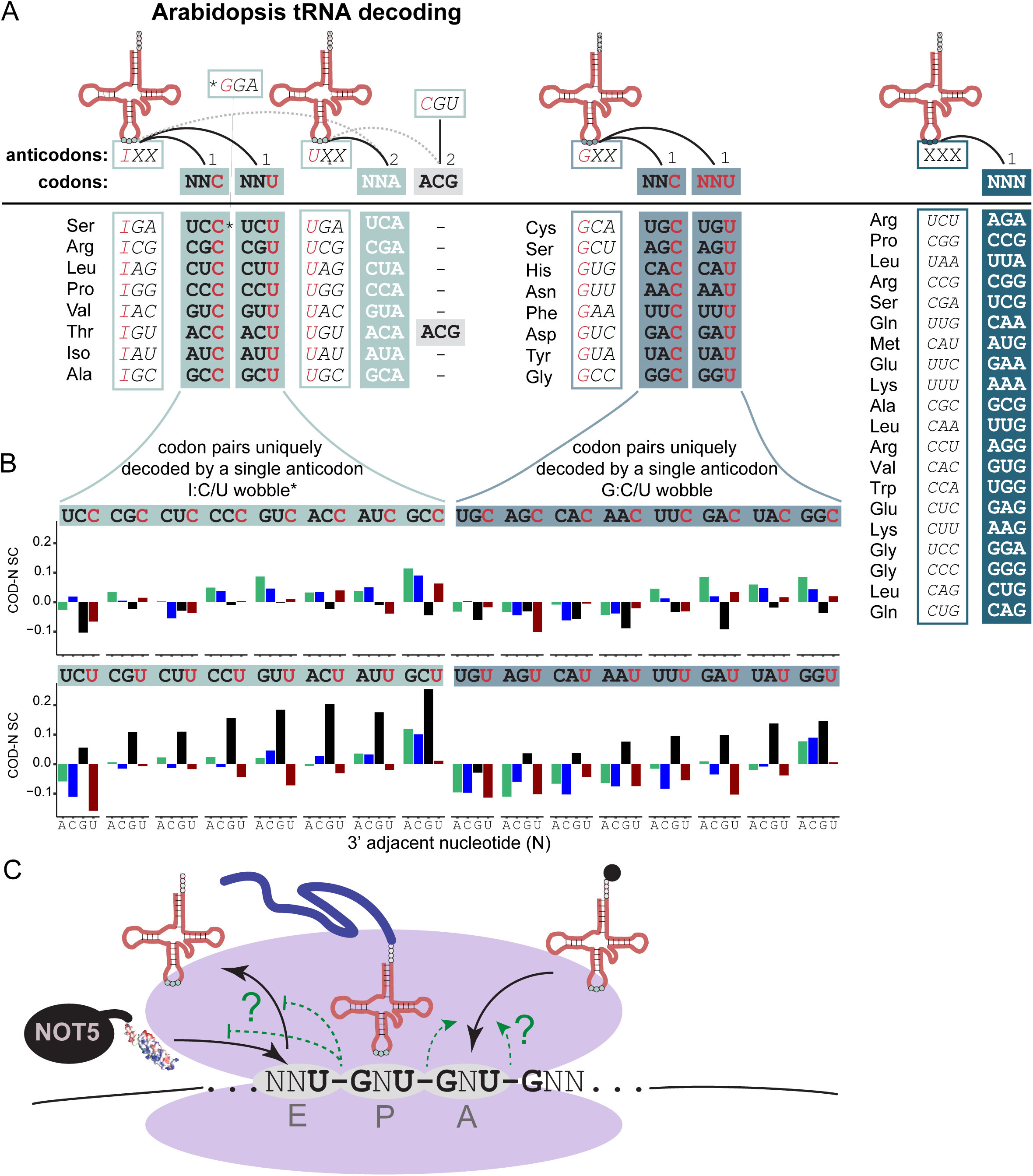
A uniquely decoded synonymous codon pair either stabilizes or destabilizes mRNAs depending on adjacent nucleotides. (A) Summary of anticodon:codon pairing and code degeneracy. Codon columns are marked with “1” if they are uniquely decoded by a single anticodon and “2” if they can be read by two different anticodons. * UCC is the exception, it is not uniquely decoded by a single anticodon, it can also be read by a GGA anticodon tRNA. (B) COD-N SC of codon pairs uniquely decoded by a single anticodon (columns: left to right, anticodons 1-8 use inosine and have I:U or I:C wobble pairing, anticodons 9-16 use G:U wobble) have different correlations depending on 3’-neighboring nucleotide (N, x-axis; A, green; C, blue; G, black; U, red). A particularly strong difference between codon pairs was observed when followed by G (black bars). (C) Adjacent nucleotide influence possible at different positions in the ribosome (UG shown). Candidate influences include A-site decoding rate, E-site tRNA release rate, and NOT5 NTD recruitment rate.

### NCANs could influence selection between codon wobble pairs

NCANs effect on mRNA stability could account for an unexpected observation in our CSC values (Figure 2A). All 16 codons pairs that only differ by U or C in their wobble positions are synonymous (e.g., Ala GCC/GCU codons). For eight of these, the wobble position is recognized by inosine in the anticodon (I:C or I:U), and for the other eight guanosine reads the wobble position (G:C or G:U). In Arabidopsis, these synonymous pairs are all uniquely decoded by the same anticodons (with one exception, Ser UCC/UCU) so, by definition, they must have the same codon optimality because the pairs use the same tRNA (Figure 5A), and yet some of these pairs have large differences in CSC values. One possibility to explain these differences is wobble decoding. Because wobble decoding occurs more slowly than a fully complementary anticodon Stadler&Fire2011), differences in tRNA-codon affinity might account for some of these CSC differences. For example, G:U and I:U wobble decoding are slower than G:C and I:C, so we would expect CSC values to be lower for the wobble codons ending in U. However, this was not the case; in four synonymous pairs the U wobble codon had a much higher CSC (e.g., GCC/GCU (Ala), AUC/AUU (Iso), CUC/CUU (Leu), and ACC/ACU (Thr)) (Figure 2A), again, suggesting that other variables were involved. NCANs can explain these differences, especially the much higher coefficients with U-G adjacent nucleotides compared with C-G (Figure 5B).

NCANs could also be a mechanism for evolutionary tuning of mRNA stability without changes to amino acid sequence or tRNA abundance. In both Arabidopsis and wheat, U-G adjacent nucleotides had correlations with RNA decay rates that were opposite of C-G suggesting opposing influence on mRNA stability. Stabilizing U-G adjacent nucleotides (e.g., N_1_N_2_U-GN_5_N_6_) could be gained or lost via a single C◊U or U◊C mutation of wobble position nucleotides next to a 3’ G (to become N_1_N_2_C-GN_5_N_6_). Similarly, when followed by a 3’ A, the opposite effect could occur, albeit less (Figure 5B). In other words, whenever one of a uniquely-decoded synonymous codon pair is used and followed by a codon that begins with G (encoding Val, Ala, Asp, Glu, or Gly) or A (Iso, Met, Thr, Asn, Lys, Ser, Arg), evolutionary tunning of mRNA decay rates could occur. Natural selection for traits favored by higher or lower mRNA stability would result in differences in frequencies of alleles only differ in synonymous codons. Critically, this selection could affect mRNA stability without changing amino acid sequence or tRNA abundance.

### Integrating codon optimality into synthetic gene encoding

Current approaches in synthetic gene design typically include a codon optimization step that involves encoding a protein based on the most used codons of the target organism. This happens because in early transgenic organisms achieving high mRNA accumulation of foreign mRNAs was surprisingly challenging [37] and coding sequence composition resulted in varying stability [38]. CUB is believed to be determined by mutation-selection equilibrium between synonymous codons [69–72]. Selection was presumed to be based on GC content or tRNA abundance. We found evidence that multiple factors influence codon usage including codon optimality and context. Other likely variables include secondary structure constraints [73]. Understanding all these influences will allow highly tailored encoding in synthetic constructs. Predictable codon influence on mRNA decay rate presents an opportunity for tailoring mRNA stability in synthetic plant genes. Estimates of codon optimality have been incorporated into models for predicting mRNA decay rate in other eukaryotes and these models guide synthetic encoding for specific mRNA stability (e.g., iCodon) [74,75]. We have confidence that this approach can now be applied to plants and suggest that consideration of NCANs might also be warranted to tailor expression.

## Conclusions

Our study addresses a gap in our understanding of codon influence on mRNA decay in plants. Codons have known associations with mRNA and protein abundance for highly expressed genes [36], so by elucidating the connection between codon frequency and mRNA decay we help to complete this connection. In doing so, we found a novel association between neighboring codons’ adjacent nucleotides and mRNA stability. This association likely reflects an unappreciated dynamic of translation that may allow variation in NOT3-mediated RNA decay rate determination through codon variation without constraining amino acid sequence.

## Methods

### Plant growth

*atnot3-1* (Salk_143332) was obtained from the Arabidopsis Biological Resource Center (https://abrc.osu.edu/). *Arabidopsis thaliana* plants were grown on growth media [0.22% (w/v) Murashige and Skoog basal salts (Caisson Labs, North Logan, UT), 1% sucrose, 1% agar (MP, Morocco), 2.3 mM 2-(4-morpholino)-ethane sulfonic acid, pH 5.7], or soil (Propagation Mix, SunGro Horticulture, Agawam, MA). Seeds were stratified in the dark at 4°C for 2 d followed by incubation in continuous cool white, fluorescent light (70 µmol photons m^-2^ s^-1^) at 22° C.

### Stabilization Coefficients and the Composite Models

In two previous studies [4,48], RNA decay estimates were generated from whole 5-d-old seedlings of *Arabidopsis thaliana* ecotype Columbia-0 (Col-0/*sov*), Col-0 expressing *proSOV:SOV*^L*er*^ (wildtype; WT), *vcs-7* (*vcs sov*), *vcs-7 proSOV:SOV*^L*er*^ (*vcs*), and *rh6 rh8 rh12 sov* after treatment with cordycepin to block transcription; these data were modeled to estimate decay rates and identify pathway substrates. For this study, we modeled decay rate of WT seedlings (independently of other genotypes) using the *RNAdecay* R package [76] and used the pathway substrate assignments of the previous studies. A decay rate value represents the probability that a specific mRNA molecule will be degraded each minute (*i.e.*, if the mRNA decay rate of a given gene = 0.01 then ∼1% of mRNAs are degraded each minute), and can be converted to the more conceptually intuitive metric of mRNA *t*_1/2_, the amount of time it takes for half of mRNAs to be degraded.

For calculating CSCs, codon frequencies were calculated as the proportion of all codons in a coding sequence excluding the start codon using the representative splice variants as designated by the Araport11 genome release [77]. Experimentally demonstrated targets of miRNAs (239 RNAs) were excluded from analysis [10]. For the CSC permutation test, scrambling CDSs and calculating trinucleotide SCs was performed programmatically in R. Other sequence unit SCs (e.g., N-COD, COD-N, N-N) were computed in a similar manner to CSCs. Decay rates were log_10_ transformed and modeled as a function of all codon frequencies or other sequence units. Least squares regression was used to estimate term coefficients in multiple linear regression models using R software. Backward selection was applied by starting with terms for all codons, non-significant terms (*p* >0.01) were removed one at a time until all remaining variables were significant. For Arabidopsis, decay rates and codon frequencies from 16,740 seedling expressed RNAs were used to generate a model predicting decay rate based uniquely on composite codon usage for any mRNA coding sequence. Alternatively, this process was repeated using only codon frequency and decay rate data from subsets of mRNAs as indicated or substitution of different sequence unit frequencies.

CSC and other sequence feature SCs from other species were calculated from frequencies in coding sequences of budding yeast (3,890 CDSs, accessed at https://yeastgenome.org), fission yeast (4,084 CDSs, https://ftp.ensemblgenomes.ebi.ac.uk/pub/fungi/release-60/fasta/schizosaccharomyces_pombe/cds/), frog (1,222 CDSs, https://www.xenbase.org/xenbase/static-xenbase/ftpDatafiles.jsp), zebrafish (3,213 CDSs, https://ftp.ensembl.org/pub/release-113/fasta/danio_rerio/cds/), human 293t cells (13,187 CDSs, https://ftp.ensembl.org/pub/current_fasta/homo_sapiens/cds/), wheat (27,794 CDSs, https://ftp.ensemblgenomes.ebi.ac.uk/pub/plants/release-60/fasta/triticum_turgidum/cds/) and compared with RNA stability measured and published by Presnyak et al. [24] (budding yeast), Eser et al. [59] (fission yeast), Bazzini et al. [26] (frog and zebrafish), Wu et al. [31] (human), and Wu et al. [3] (wheat).

### Transgene construction and transformation

The following were purchased as cloned synthetic DNA fragments (ThermoFisher Scientific): the attL1 recombination sequence fused to 634 bp genomic sequence upstream of the Arabidopsis *UBQ10* start codon [49] fused to a SpeI restriction site, YFP sequences encoded for maximal and minimal predicted stability (Supplemental Data 17) fused to a 5’ SpeI restriction site and to the attL2 recombination sequence 3’ of the YFP sequence. SpeI restriction sites were used to fuse the *UBQ10* promoter to YFP variants and recombination sequences were used to clone the gene constructs into the plant destination vector pGWB501 [78] using Gateway™ LR Clonase™ II Enzyme mix (Thermo Fisher Scientific). Recoded *DJC23* (AT4G36040) CDSs were purchased from ThermoFisher Scientific and cloned into the pENTR vector (Life Technologies). Constructs were made using a multi-stie recombination reaction (LR Clonase™ II) of pCAM-kan-R4R3 binary destination vector, p1R4-ML-XVE (supplied by the University of Helsinki), pENTR-clones, and p2R3a-nosT (supplied by Addgene) according to Siligato et al. [51]. Gene sequences were verified by Sanger sequencing at the University of Utah Genomics Core Facility. Transgenic plant lines were generated using the resulting gene construct-containing binary vectors using *Agrobacterium tumefaciens (GV3101-pMP90)*-mediated floral dip transformation [79]. Transformants were identified by resistance to hygromycin B (15 µg/mL) or kanamycin (50 µg/mL) on growth media. Lines containing a single locus insertion were identified by segregation analysis of hygromycin resistance.

### Alignment

For alignment of the Not3 N-terminal domain, yeast Not5 orthologous sequences were found on NCBI (https://www.ncbi.nlm.nih.gov): QHB12322.1 NOT5_YEAST, ref|XP_717644.2| CCR4-NOT core subunit [Candida albicans SC5314], ref|XP_005849464.1| hypothetical protein CHLNCDRAFT_142739 [Chlorella variabilis], ref|NP_001330084.1| transcription regulator NOT2/NOT3/NOT5 family protein [Arabidopsis thaliana], ref|XP_020402447.1| CCR4-NOT transcription complex subunit 3 isoform X1 [Zea mays], ref|XP_003624606.1| CCR4-NOT transcription complex subunit 3 isoform X1 [Medicago truncatula], ref|XP_065514074.1| CCR4-NOT transcription complex subunit 3 [Caloenas nicobarica], ref|XP_015269889.1| CCR4-NOT transcription complex subunit 3 [Gekko japonicus], ref|NP_001315047.1| CCR4-NOT transcription complex subunit 3a [Danio rerio], ref|XP_018082443.1| CCR4-NOT transcription complex subunit 3 S homeolog isoform X1 [Xenopus laevis], ref|NP_666288.1| CCR4-NOT transcription complex subunit 3 [Mus musculus], ref|NP_055331.1| CCR4-NOT transcription complex subunit 3 [Homo sapiens]. ClustalW was used to align sequences [80].

### tRNA analysis

Folding predictions and isotype analysis for 206 unique tRNA sequences from 661 genes were made using tRNAscan-SE2 [81] (Supplemental Data 1) to identify conservation of NOT3 interaction residues. In the Arabidopsis thaliana genome, 661 complete tRNA genes with clear isotype-anticodon agreement (2 mitochondrial, 36 plastid) contain 206 unique nuclear-encoded tRNA sequences with 74 unique folding structures predicted.

### YFP fluorescence

Root tip images were captured on an Olympus BX50 microscope using an Olympus DP71 camera. YFP fluorescence in root tips was captured under Hg lamp illumination using the Olympus U-MWIBA filter cube (excitation 460–490 nm; emission 515–550 nm). 16 bit images were captured using 3 s exposure time and ISO 200. Root tip YFP intensity was reported as mean pixel intensity of a circular area inscribed in the root apex region.

### RNA decay analysis

RNA decay analysis was performed as described previously (Sorenson et al., 2018a). Briefly, seedlings were pre-incubated in incubation buffer (15 mM sucrose, 1 mM Pipes pH 6.25, 1 mM KCl, 1 mM sodium citrate) with shaking 125 rev min^-1^ aeration for 15 min. Untreated samples were collected (T_0_) or buffer was exchanged with fresh buffer containing 1 mM cordycepin and incubated for an additional 15, 60, or 240 min and flash frozen. RNA was extracted from pulverized flash-frozen whole seedlings using the Quick-RNA Plant Miniprep Kit (Zymo Research, Irvine, CA). RNA was treated with DNase I (RNase-Free) (New England Biolabs, Ipswich, Massachusetts). 2 µg RNA were reverse transcribed using random hexamers for priming and the High-Capacity cDNA Reverse Transcription Kit (Thermo Fisher Scientific). Quantitative reverse transcription polymerase chain reaction was used to evaluate cDNA abundances using PowerUp™ SYBR® Green Master Mix (Thermo Fisher Scientific) and the CFX96 Real-Time Detection System (Bio-Rad). The following stable reference genes (each with a *t*_½_ > 48 h) were used to normalize abundance across the decay experiment time course according to the mathematical framework of Hellemans et al. [82]: *UBC10* (At5g53300), *VHA-A* (At1g78900), and *eEF1B* (At1g57720) for YFP^stCOD^ and YFP^unCOD^. For *DJC23^stCod^* and *DJC23^unCod^ UBC10* was used as the reference RNA. Primer sequences are listed in Supplemental Data 18. RNA decay rates were estimated using our previously described modeling approach [4,76].

### Feature and Gene Ontology Analysis

For mRNA feature analysis, CDS length, 5’-UTR length, and 3’UTR length were determined from the TAIR10 genome annotation release [83]. Lengths were of the designated representative splice variants. Feature lengths were binned into 40 quantiles and mean lengths of each bin were reported.

For GO analysis, annotations were downloaded from https://www.arabidopsis.org as the “ATH_GO_SLIM.txt” file from the data release dated 12-31-2024. GO term definitions were downloaded from https://geneontology.org as the “go-basic.obo” file on February 18, 2025. Enrichments were tested using the *topGO* R package from https://bioconductor.org, applying the “weight01” algorithm to perform Fisher’s exact test, followed by the method of Benjamini and Hochberg [84] to control the false discovery rate.

## Declarations

### Consent for publication

Not applicable.

### Availability of data and materials

Data analyzed in this study are found in the Gene Expression Omnibus (http://www.ncbi.nlm.nih.gov/geo) under accession numbers GSE86361 and GSE136713. Fit model parameters are included in this published article and its supplementary information files.

### Authors’ contributions

R.S.S., and L.S. conceived and designed experiments, analyzed data, and wrote the manuscript; R.S.S. performed experiments, performed computation.

## Acknowledgements

We would like to thank Dr. Fred Adler, Katrina Johnson, Kelly Hughes, and Fabienne Chevance for helpful discussions.

## Additional Files

Additional File 1.xlsx – 18 worksheets:

1. Supplemental Data 1. Table of linear regression values. Log10 decay rate was regressed on -log2 relative codon bias (relative to genome-wide codon bias) for each codon separately. Significant slopes (p value < 0.01) are red.
2. Supplemental Data 2. Trinucleotide SCs from scrambled CDSs (x10,000), codons, +1 frame, +2 frame.
3. Supplemental Data 3. Table of multiple regression values from regression of decay rate on composite codon frequency. Decay rate (log10) was modeled as the sum of codon terms and a constant. Codon terms were composed of the product of codon frequency variable and its coefficient. Coefficients were estimated through multiple regression on 16,935 Arabidopsis seedling mRNAs and indicate how much influence each codon frequency has on decay rate.
4. Supplemental Data 4. Codon decay coefficients of codons with significant terms from multiple regression of log10 decay rate on composite codon frequency using 7,555 mRNAs that are more than 50% VCS-dependent in a Col-0/sov background.
5. Supplemental Data 5. Codon decay coefficients of codons with significant terms from multiple regression of log10 decay rate on composite codon frequency using 3445 mRNAs that are more than 50% VCS-dependent in an SOV background.
6. Supplemental Data 5. Codon decay coefficients of codons with significant terms from multiple regression of log10 decay rate on composite codon frequency using 6020 mRNAs that are more than 50% RH-dependent in a Col-0/sov background.
7. Supplemental Data 7. Codon decay coefficients of codons with significant terms from multiple regression of log10 decay rate on composite codon frequency using 6007 mRNAs that are less than 50% RH&VCS (with or without SOV) dependent.
8. Supplemental Data 8. tRNAscan-SE2_folding results for 206 unique tRNA sequences. After folding prediction, sequences were checked for conservation of NOT3 interacting residues.
9. Supplemental Data 9. GO enrichment among genes with the 5 pct highest Arg codons AGA and CGG combined frequency.
10. Supplemental Data 10. Contiguous dicodon SCs. Compare Figure 4A.
11. Supplemental Data 11. Reciprocal dicodon SC differences (ΔdiCSC).
12. Supplemental Data 12. Neighboring codon adjacent nucleotides SCs.
13. Supplemental Data 13. Codon + 5’-nt (N-COD) and codon + 3’-nt (COD-N) SCs.
14. Supplemental Data 14. Composite COD-N multiple regression model coefficients for Arabidopsis. Table of multiple regression values from regression of decay rate on composite COD-N frequency. Decay rate (log10) was modeled as the sum of COD-N terms and a constant. Terms were composed of the product of COD-N frequency and its coefficient. Coefficients were estimated through multiple regression on 16,935 Arabidopsis seedling mRNAs and indicate how much influence each COD-N frequency has on decay rate.
15. Supplemental Data 15. Table of multiple regression values from regression of decay rate on composite codon frequency. Decay rate (log10) was modeled as the sum of codon terms and a constant. Codon terms were composed of the product of codon frequency variable and its coefficient. Coefficients were estimated through multiple regression on 33,430 wheat mRNAs and indicate how much influence each codon frequency has on decay rate (Wu et al. (2024).
16. Supplemental Data 16. Composite COD-N mupltiple regression model coeficients for wheat. Table of multiple regression values from regression of decay rate on composite COD-N frequency. Decay rate (log10) was modeled as the sum of COD-N terms and a constant. Terms were composed of the product of a COD-N frequency variable and its coefficient. Coefficients were estimated through multiple regression on 33,430 wheat mRNAs and indicate how much influence each codon frequency has on decay rate (Wu et al. (2024).
17. Supplemental Data 17. Coding sequences of YFP synonymously encoded with codons used by stable mRNAs (YFP^stCod^) or unstable mRNAs (YFP^unCod^) based on mRNA decay rate associations with codon usage.
18. Supplemental Data 18. Table of primers used in this study.

## Abbreviations

CUB: codon usage bias
ORF: open reading frame
CDS: coding sequence
COMD: codon optimality-mediated decay
CSC: codon stabilization coefficient
SC: stabilization coefficient
UTR: untranslated region
GO: gene ontology
YFP: yellow fluorescent protein
NTD: N-terminal domain
ΔdiCSC: reciprocal dicodon stabilization coefficient difference
dicodon NCAN: neighboring codon adjacent nucleotides.

---

**Supplemental Figure 1.**
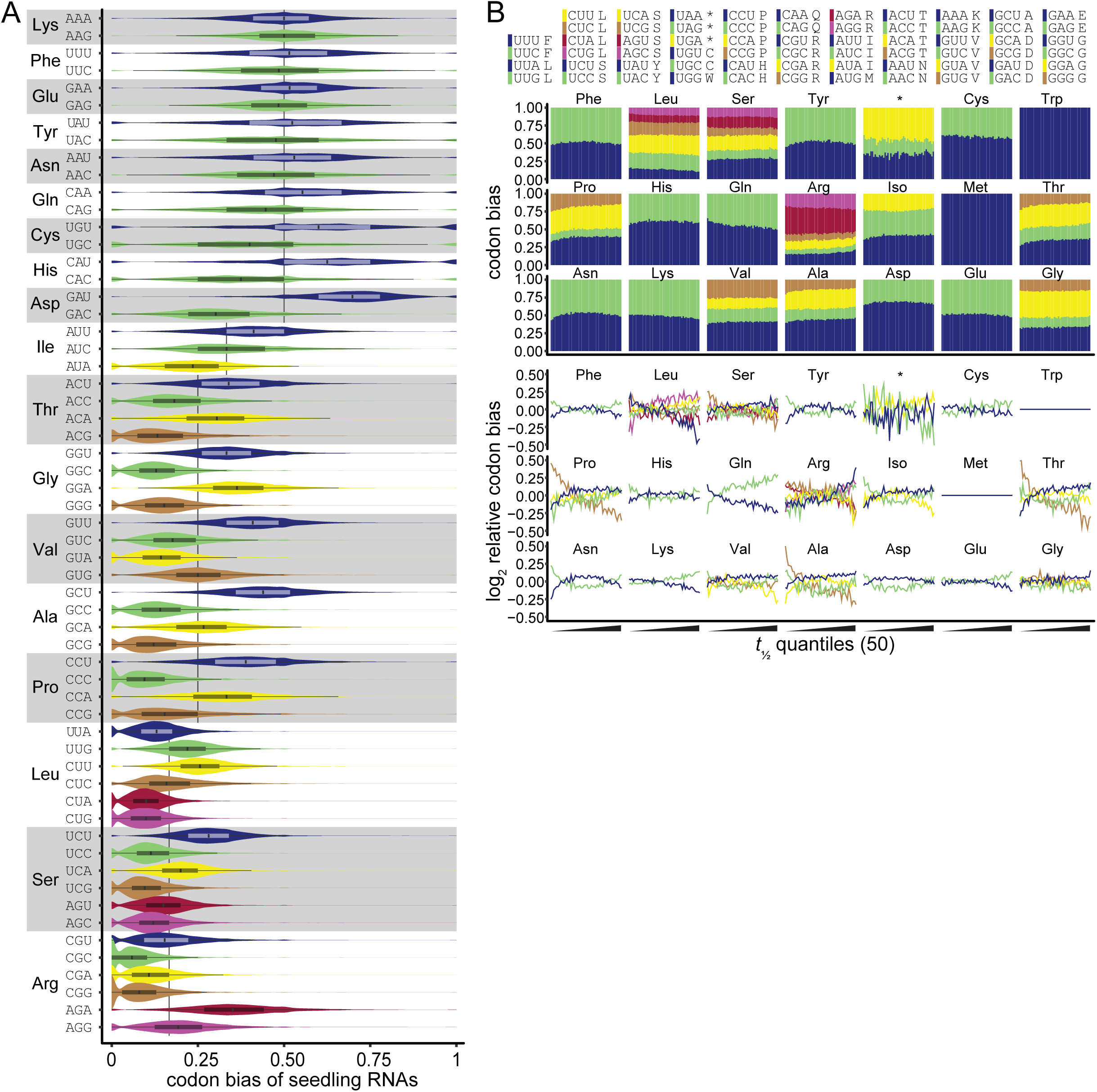
Individual mRNAs vary in codon usage. (A) Distribution of codon biases of 16,749 individual transcripts expressed in 5-d-old Arabidopsis thaliana seedlings. Codon bias was calculated for each transcript; if a transcript did not encode a given amino acid it was not counted in the distribution. Boxplots are overlaid on violin plots showing codon bias distributions. Vertical black lines behind violin plots represent expectation if codon usage for a given amino acid was random. Note: Trp and Met have a single codon, therefore do not have bias, but are included for completeness. (B) Codon bias (upper) and log2 codon bias relative to genome-wide codon bias for mRNA *t*_1/2_ quantiles (increasing left to right; subset of data in Figure 1C).

**Supplemental Figure 2.**
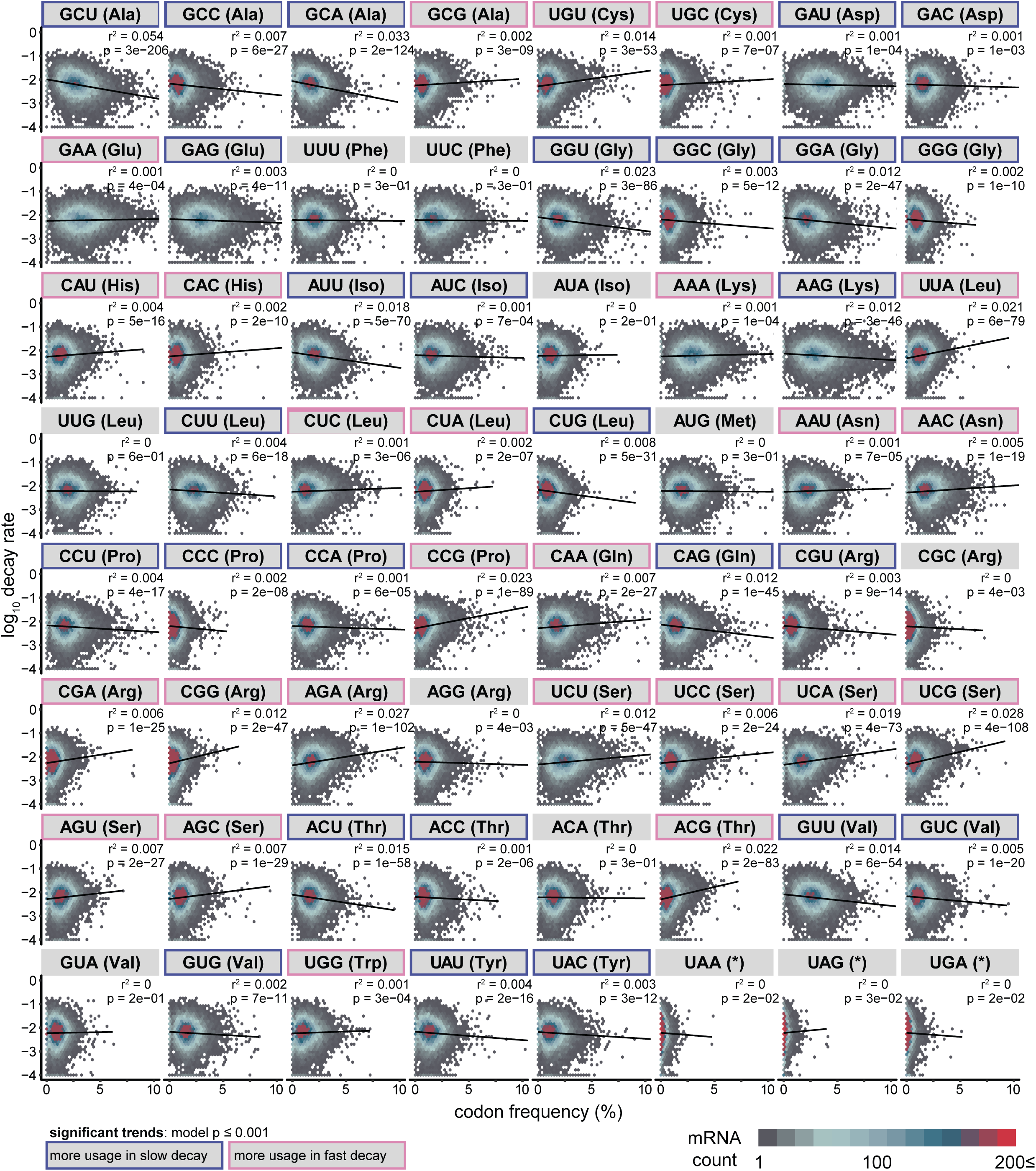
Codon frequencies correlate with mRNA *t*_1/2_. A scatter-density plot for each codon compares codon frequency (as percent) with log10 mRNA decay rates. Linear regression fit is indicated by black lines and r2 and p-values label each plot. Hexagon colors indicate point (individual mRNA) density. Codons are presented alphabetically by amino acids and significant trends are highlighted by outline.

**Supplemental Figure 3.**
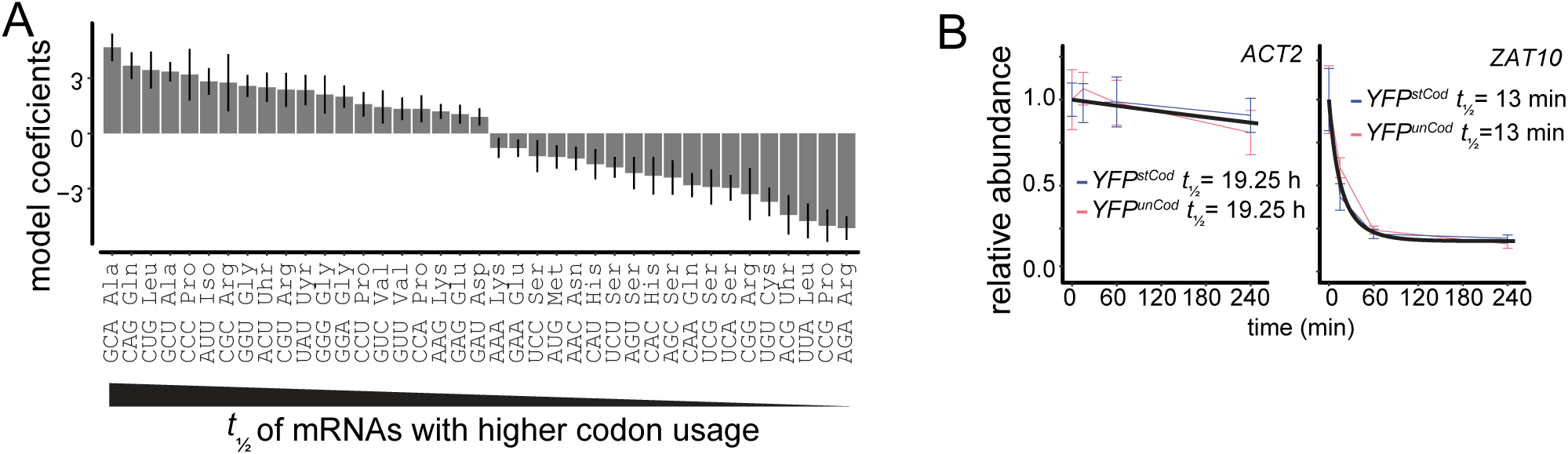
Transcript features and codon usage. (A) Multiple regression model coefficients for significant codons indicates the size of the influence of each codon on decay rate. (B) ACT2 and ZAT10 *t*_1/2_s from *proUBQ10::YFPstCOD* and *proUBQ10::YFPunCOD* transgenic plants.

**Supplemental Figure 4.**
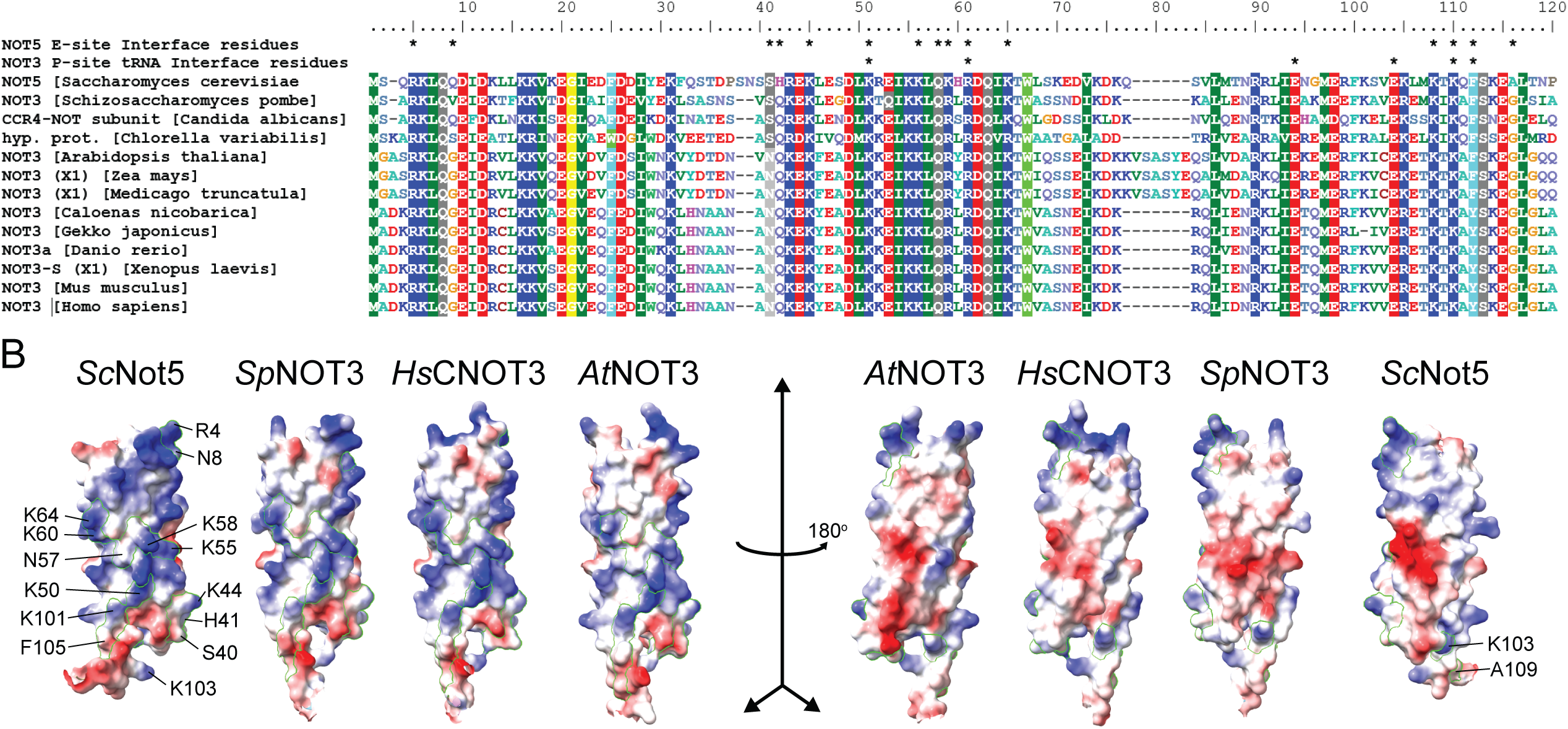
AtNOT3 NTD conservation. (A) Alignment of yeast Not5 N-terminal region from representative eukaryotes homologs. Residues shown to interact with the ribosome E-site by Buschauer et al. (2020) or P-site tRNA by Zhu et al. (2024) are indicated with asterisks. Comparing Sc with At directly, identical residues: R4, K44, K50, K55, Q57, R60, K64, K101, K103, F105; similar residues: Sc->At: K58->R, A109->G, different and conserved in all other species (Sc->At): Q8->G, S40->N, H41->Q. (B) NTD sensor structure (AlphaFold) comparison with ScNot5 E-site interface residues labeled. Electrostatic charge is colored as red (positive), white (neutral), and blue (negative) [54].

**Supplemental Figure 5.**
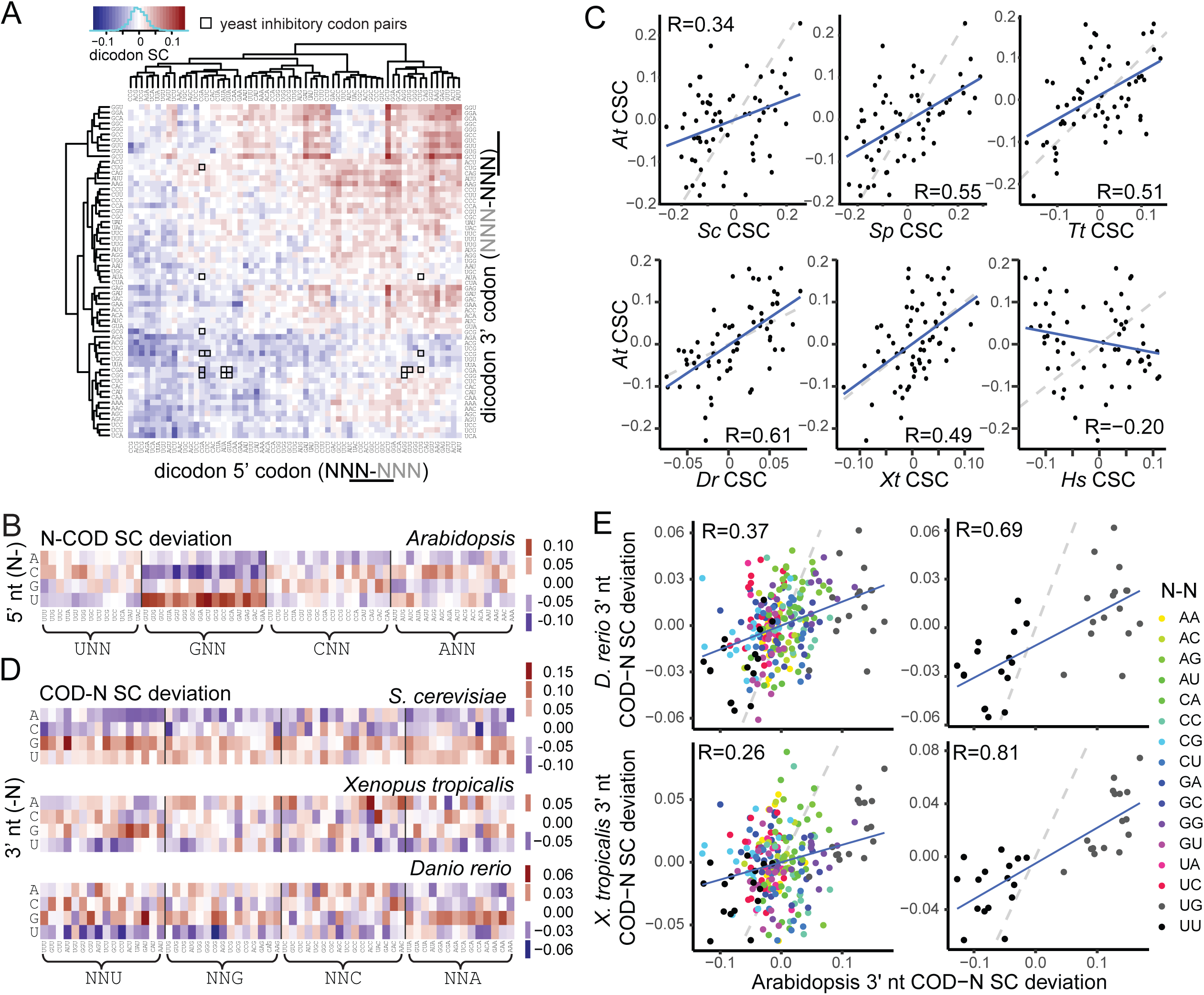
SCs calculated for frameshifted trinucleotides suggest neighboring codons contribute to codon-mediated stability variation. (A) SCs of dicodons. Hierarchical clustering of codons in the first (5’) or second (3’) position of the dicodon. Outlined boxes indicate SCs 17 dicodons (AGG-CGA, AGG-CGG, AUA-CGA, AUA-CGG, CGA-AUA, CGA-CCG, CGA-CGA, CGA-CGG, CGA-CUG, CGA-GCG, CUC-CCG, CUG-AUA,, CUG-CCG, CUG-CGA, GUA-CCG, GUA-CGA, GUG-CGA) found to be inhibitory to translation in a wobble-decoding dependent manner in yeast by Gamble et al. (2016). (B) The effect of a the 5’-neighboring nucleotide on codon SC. Heatmap of N-COD SC deviation (mean of N-COD SC by codon - N-COD SC). (C) CSC comparison of Arabidopsis with S. cerevisiae, S. pombe, X. tropicalis, D. rerio, T. turgidum, and H. sapiens. Plots are labeled with Pearson correlation values. (D) The effect of a the 3’-neighboring nucleotide on codon SC. Heatmap of COD-N SC deviation (mean COD-N SC by codon - COD-N SC) for S. cerevisiae, X. tropicalis and D. rerio. (E) Comparison of Arabidopsis COD-N SC deviation with those of X. tropicalis and D. rerio. COD-N combinations are colored by the codon 3’ nucleotide and the adjacent nucleotide pair. All COD-N combinations (top) and just those with UG or UU adjacent nucleotides (bottom).

